# Auditory attention alterations in migraine: a behavioral and MEG/EEG study

**DOI:** 10.1101/661413

**Authors:** Rémy Masson, Yohana Lévêque, Geneviève Demarquay, Hesham ElShafei, Lesly Fornoni, Françoise Lecaignard, Dominique Morlet, Aurélie Bidet-Caulet, Anne Caclin

**Affiliations:** Lyon Neuroscience Research Center (CRNL), INSERM UMRS 1028, CNRS UMR 5292, Université Claude Bernard Lyon 1, Université de Lyon, Lyon, France; Neurological Hospital Pierre Wertheimer, Functional Neurology and Epilepsy Department, Hospices Civils de Lyon and Université Claude Bernard Lyon 1, Université de Lyon, Lyon, France

**Keywords:** Migraine, event-related responses, bottom-up attention, top-down attention, electroencephalography, magnetoencephalography

## Abstract

**Objectives:** To evaluate alterations of top-down and/or bottom-up attention in migraine and their cortical underpinnings.

**Methods:** 19 migraineurs between attacks and 19 matched control participants performed a task evaluating jointly top-down and bottom-up attention, using visually-cued target sounds and unexpected task-irrelevant distracting sounds. Behavioral responses and MEG/EEG were recorded. Event-related potentials and fields (ERPs/ERFs) were processed and source reconstruction was applied to ERFs.

**Results:** At the behavioral level, neither top-down nor bottom-up attentional processes appeared to be altered in migraine. However, migraineurs presented heightened evoked responses following distracting sounds (orienting component of the N1 and Re-Orienting Negativity, RON) and following target sounds (orienting component of the N1), concomitant to an increased recruitment of the right temporo-parietal junction. They also displayed an increased effect of the cue informational value on target processing resulting in the elicitation of a negative difference (Nd).

**Conclusions:** Migraineurs appear to display increased bottom-up orienting response to all incoming sounds, and an enhanced recruitment of top-down attention.

**Significance:** The interictal state in migraine is characterized by an exacerbation of the orienting response to attended and unattended sounds. These attentional alterations might participate to the peculiar vulnerability of the migraine brain to all incoming stimuli.

**Highlights:**

- Migraineurs performed as well as healthy participants in an attention task.
- However, EEG markers of both bottom-up and top-down attention are increased.
- Migraine is also associated with a facilitated recruitment of the right temporo-parietal junction.

## 1. Introduction

Migraine is the most common neurological disorder with a prevalence around 10% in the worldwide population (Stovner et al., 2007). Migraine is mainly characterized by recurrent headache attacks often accompanied by nausea and vomiting, all of which can be disabling and have a vast impact on quality of life. Migraine attacks are strongly associated with photophobia, phonophobia, osmophobia (aversion to visual, auditory and olfactory stimuli, respectively), and allodynia (pain sensitization to non-painful somatosensory stimuli) (Headache Classification Committee of the International Headache Society (IHS), 2013). These “phobias” encompass both a heightened sensitivity to external stimulation and an exacerbation of pain by those same stimulations. Sensory alterations persist, to a smaller extent, during the attack-free period. Interictally, thresholds for light-induced discomfort or pain (Main et al., 1997; Vanagaite et al., 1997) were found decreased in migraine (i.e., hypersensitivity), and intensity of light-induced pain was found exacerbated (Drummond, 1986). Similar results were reported in the auditory modality (Main et al., 1997; Vingen et al., 1998) and migraineurs describe a general over-responsiveness to everyday non-noxious stimuli in subjective questionnaires (Granovsky et al., 2018; Lévêque et al., 2019).

EEG is a particularly useful technique to investigate sensory processing with its high temporal resolution which allows a fine understanding of transient responses to sensory stimulation (Schoenen et al., 2003). Regarding the interictal period, the main result reported by previous EEG studies was a lack of habituation of brain responses to repeated visual stimulation (for a review, see Coppola et al., 2009). Deficits of habituation in migraine were described for various event-related potentials (ERPs): mostly for sensory components such as the visual P1 and N1 (Áfra et al., 2000; Ozkul and Bozlar, 2002; Schoenen et al., 1995), but also for later cognitive ERPs such as the P3b (Evers et al., 1999; Siniatchkin et al., 2003) and the contingent negative variation (CNV) (Kropp et al., 2015; Kropp and Gerber, 1993; Schoenen and Timsit-Berthier, 1993). Interestingly, those habituation impairments normalize before and during migraine attacks (Evers et al., 1999; Judit et al., 2000; Kropp and Gerber, 1995), even though hypersensitivity climaxes during attacks. Impairment of habituation in migraineurs is considered a hallmark of migraine neurophysiology and a biomarker of the interictal state in migraine. However, these results have not been replicated in recent studies (Omland et al., 2016, 2013; Sand and Vingen, 2000). In the auditory modality, studies investigating habituation deficits in migraine are much scarcer and produced negative results (Morlet et al., 2014; Sand and Vingen, 2000; Wang and Schoenen, 1998). Other EEG responses have also been investigated, notably steady-state visual evoked potentials (SSVEP), electrical brain responses to repeated visual stimulations at specific frequencies. Results strongly suggest that the excitability of the occipital cortex is abnormal among migraineurs (de Tommaso, 2019; de Tommaso et al., 2014).

Magnetoencephalography (MEG) is also a powerful tool for the investigation of sensory processing. It provides a superior signal-to-noise ratio which allows for precise source reconstruction and a better sensitivity to sources tangential to the scalp. In addition, MEG studies of sensory processing migraine are still very scarce because of the few available MEG systems and experts in the world. Nevertheless, the few existing MEG studies appear to confirm results obtained using EEG (Chen et al., 2013; Korostenskaja et al., 2011). Further investigating migraine using MEG could provide new insights in migraine pathophysiology.

It is still unclear if the sensory dysfunction in migraine is only rooted in alterations of “low-level” stages of sensory processing or if the impairment of cognitive processing of sensory inputs also plays a part. It has been established that poor cognitive performance is associated with migraine attacks and sometimes persists during the interictal period (Vuralli et al., 2018). During a passive auditory oddball task, enhanced amplitudes of the N1 orienting component (Morlet et al., 2014) and of the P3a (Demarquay et al., 2011) have been reported among migraineurs. These two ERPs have been associated with the involuntary orienting of attention (Näätänen and Picton, 1987; Polich, 2007). In the visual modality, migraineurs were also found to present a heightened involuntary attentional orienting, a decreased ability to suppress unattended stimuli in the periphery, and abnormalities in top-down attentional processes (Mickleborough et al., 2011a). This is corroborated by reports of self-perceived attentional difficulties by migraineurs (Carpenet et al., 2019; Lévêque et al., 2020; Sacks, 1992). Furthermore, some clinic-based studies using neurophysiological tests revealed that migraine had a moderate effect on attentional performances during the interictal period (reviewed in Vuralli et al., 2018). However, attention impairment was not consistently detected in clinical studies (Burker et al., 1989; Conlon and Humphreys, 2001; Koppen et al., 2011) and the precise attentional mechanisms altered in migraine remain to be characterized.

The present study aims to better characterize which attentional brain mechanism is potentially impaired in migraine. In a world saturated with sensory information, the allocation of our limited cognitive processing resources is guided by two main attentional processes. Top-down (or voluntary) attention enables to selectively attend stimuli which are relevant to our goals, and to filter out irrelevant stimuli. It operates through inhibitory and anticipatory mechanisms (Bidet-Caulet et al., 2010), underpinned by the dorsal attention network (Corbetta et al., 2000; Corbetta and Shulman, 2002) and reflected in EEG by specific ERPs such as the Contingent Negative Variation (CNV, (Brunia and van Boxtel, 2001) or the Negative Difference (Nd, (Alcaini et al., 1994a; Giard et al., 2000; Näätänen, 1982). As for bottom-up (or involuntary) attention, it is the ability to have our attention captured by unexpected salient events in one’s environment. It is mediated by the ventral attention network (Corbetta et al., 2008; Corbetta and Shulman, 2002) and reflected in EEG by the ERPs such as the orienting component of the N1 and the P3a (orienting of the attention towards the unexpected stimulus, see Alcaini et al., 1994b; Escera et al., 2000; Simons et al., 2001; Yago et al., 2003) and the reorienting negativity (RON, reorienting of the attention back to the task at hand, see Munka and Berti, 2006; E. Schröger and Wolff, 1998). Based on previous studies, we hypothesize that migraine is associated with exacerbated bottom-up and/or deficient top-down attention processes, resulting in the inability to filter out irrelevant information and possibly participating to the sensory disturbances associated with this disorder. To this day, very few electrophysiological studies (see above) have attempted to investigate attention in migraine.

Migraineurs and control participants were recruited to perform an adapted version of the Competitive Attention Task (Bidet-Caulet et al., 2015) while brain activity was monitored using EEG and MEG. This paradigm enables to conjointly evaluate top-down and bottom-up attention, using visually-cued target sounds and unexpected task-irrelevant distracting sounds. The Competitive Attention Task has been successful in investigating specifically both facets of attention in healthy young adults (Bidet-Caulet et al., 2015; ElShafei et al., 2018a, 2019; Masson and Bidet-Caulet, 2019), in children (Hoyer et al., 2019) and in the elderly population (ElShafei et al., 2018b). Analyses of behavioral performances, event-related potentials, and event-related fields both at the sensor and source levels were conducted to detect any attention alterations in migraine.

## 2. Materials and methods

### 2.1. Participants

25 migraine patients (17 female, 8 male) suffering from migraine without aura were included in this study. Inclusion criteria were age between 18 and 60 years (Table 1) and have a diagnosis of migraine with a reported migraine frequency between 2 to 5 days per month. Exclusion criteria comprised migraine with aura, chronic migraine, and migraine preventive medication. Every patient was examined by a neurologist (GD, Hospices Civils de Lyon). Migraine patients filled out the Hospital Depression and Anxiety scale (Zigmond and Snaith, 1983), the HIT-6, a short questionnaire aiming to evaluate headache impact on everyday life (Kosinski et al., 2003) and the Migraine Disability Assessment Questionnaire (MIDAS) (Stewart et al., 1999). Several EEG studies have reported a normalization of electrophysiological markers of the interictal period of migraine during the peri-ictal period (just before and just after migraine attacks), a normalization which peaks during the migraine attack (Chen et al., 2009; Evers et al., 1999; Judit et al., 2000; Kropp and Gerber, 1995; Mulder et al., 1999). As we were interested in studying attention during the interictal state, if the patient had a migraine attack during the 72 hours before the testing session, the session was postponed to an ulterior date. If the patient had a migraine attack during the 72 hours after the session, collected data were not used in the analyses, as it is common practice in neuroimaging studies of migraine (Demarquay and Mauguière, 2016). Data from 19 patients (13 female, 6 male) were usable in this study: data from 5 patients were discarded because a migraine attack happened in the 72 hours following the recording session and data from 1 patient because the patient failed to perform the task correctly.

**Table 1:** Demographics and headache profile of the control and migraine groups. Two control participants did not filled the Hospital Anxiety and Depression (HAD) scale. Mean and standard deviation are provided. Group differences are tested using a non-parametric Mann-Whitney U test. NA: not applicable.

|  | **Migraine** | **Control** | **p-value** |
| --- | --- | --- | --- |
| **Sample size** | 19 | 19 | - |
| **Age (years)** | 32.7 (8.7) | 31.2 (7.8) | 0.53 |
| **Sex (number of female participants)** | 13 (68%) | 13 (68%) | - |
| **Education level (years)** | 15.8 (3.1) | 15.8 (2.2) | 0.99 |
| **Musical practice (years)** | 2.8 (3.3) | 2.8 (3.5) | 0.74 |
| **Laterality (number of right-handed)** | 19 | 19 | - |
| **Anxiety score** | 5.7 (3.5) | 4.6 (2.5) | 0.42 |
| **Depression score** | 2.6 (2.6) | 1.8 (2.0) | 0.31 |
| **Migraine duration (years)** | 16.8 (7.4) | NA | - |
| **HIT-6 score** | 64.2 (7.1) | NA | - |
| **MIDAS score** | 12.8 (12.1) | NA | - |

19 control participants free of migraine and matched to the patients for sex, age, laterality, education level, and musical practice^1^ were included in this study. Exclusion criteria for all subjects included a medical history of psychological or neurological disorders, ongoing background medical treatment other than contraceptive medication, pregnancy, and hearing disability. All subjects gave written informed consent and received a monetary compensation for their participation.

### 2.2. Task and procedure

75 % of the trials consisted in a visual cue (200 ms duration) followed after a 1000 ms delay by an auditory target (100 ms duration with 5 ms rise-time and 5 ms fall-time) (Figure 1a). The cue was centrally presented on a screen (gray background) and could be a green arrow pointing to the left, to the right, or to both sides. The target sounds were monaural pure sounds presented at 25 dB SL. The low-pitched target sound had a fundamental frequency of 512 Hz, the high-pitched target was 2 semi-tones higher than the low-pitched sound (574 Hz). If during training the subject was unable to discriminate the two sounds, the pitch difference could be increased up to 3 semi-tones by steps of half a semi-tone prior to starting EEG/MEG recordings.

**Figure 1:**
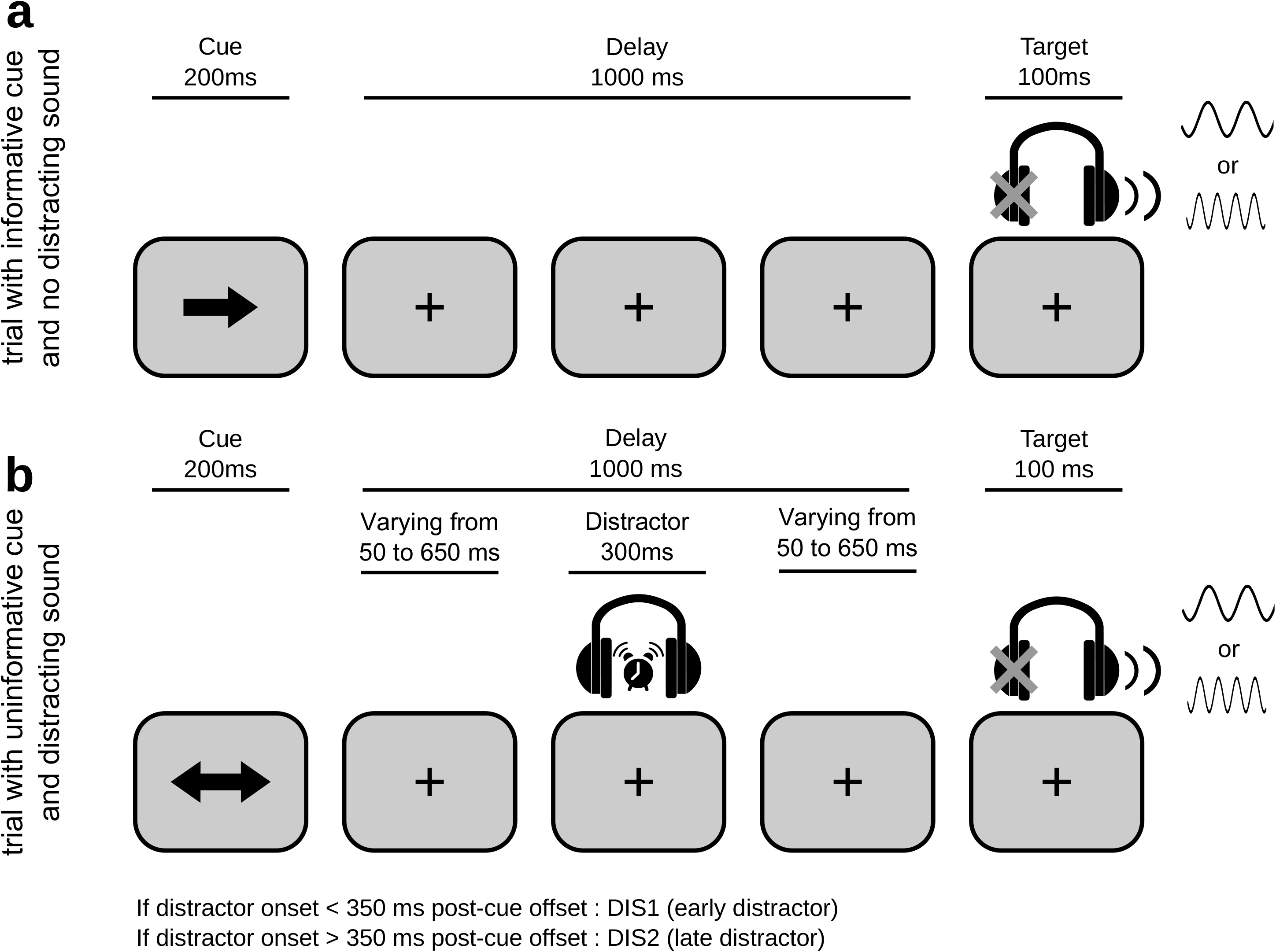
Protocol. The task was to discriminate between a low- and a high-pitched sound, presented monaurally. A visual cue initiated the trial, and was either informative (50%) or non-informative (50%) about the target ear. 25% of the trials included a distracting sound. **(a)** Example of an *informative* trial with no distracting sound: a one-sided visual cue (200 ms duration) indicates in which ear (left or right) the target sound (100 ms duration) will be played after a fixed 1000 ms delay. **(b)** Example of an *uninformative* trial with a distracting sound: a two-sided visual cue (200 ms duration) does not provide any indication in which ear (left or right) the target sound will be played. The target sound can be a high- or low-pitched sound indifferently of the cue informational value. In 25% of all trials (with *informative* or *uninformative* cues), a loud binaural distracting sound (300 ms duration), such as a clock ring, is played during the cue-target interval at a random delay after the cue offset: the *DIS1* condition corresponds to early distracting sounds (starting 50–350 ms after cue offset), the *DIS2* condition corresponds to late distracting sounds (starting 350–650 ms after cue offset).

In the other 25 %, the same trial structure was used, but a binaural distracting sound (300 ms duration, 55 dB SL) was played at some point between the cue offset and the target onset (Fig.1b). Trials with a distracting sound starting between 50 ms and 350 ms after the cue offset were classified as *DIS1* (early distracting sound), those with a distracting sound starting between 350 ms and 650 ms after the cue offset were classified as *DIS2* (late distracting sound), and those with no distracting sound were classified as *NoDIS*. A total of 40 different ringing sounds were used as distracting sounds (clock-alarm, door-bell, phone ring, etc.) in each participant.

The cue and target categories (i.e., *informative* vs. *uninformative* cue, low vs. high-pitched tone) were presented in the same proportion for trials with and without distracting sound. In 25 % of the trials, the cue was indicating left and the target was presented on the left side. In 25 % of the trials, the cue was indicating right and the target was presented on the right side. This leads to a total of 50 % of *informative* trials. In the last 50 % of the trials, the cue was *uninformative*, indicating both directions and the target was presented equiprobably on the left or right side. The target type (high or low) was presented in the same proportion (50% each) in all conditions. To compare brain responses to acoustically matched sounds, the same distracting sounds were played in each combination of cue category (*informative*, *uninformative*) and distractor condition (*DIS1* or *DIS2*). Each distracting sound was thus played 4 times during the whole experiment, but no more than once during each single block to limit habituation.

Participants were instructed to perform a discrimination task and to respond as fast as possible by pushing or pulling a joystick. The mapping between the targets (low or high) and the responses (pull or push) was counterbalanced across participants, but did not change across the blocks, for each participant. They were asked to allocate their attention to the cued side in the case of informative cues. Participants were informed that informative cues were 100 % predictive and that a distracting sound could be sometimes played. Participants had a 3.4 second response window. At any time in the absence of the visual cue, a blue fixation cross was presented at the center of the screen. Participants were instructed to keep their eyes fixating on the cross and to minimize eye movements while performing the task.

Participants were in a seating position. All stimuli were delivered using Presentation software (Neurobehavioral Systems). Auditory stimuli were delivered through air-conducting plastic ear tubes. First, the auditory threshold was determined for the low-pitched target sound, in each ear, for each participant using the Bekesy tracking method (Leek, 2001). Second, participants were trained with a short sequence of the task (task difficulty was adjusted if needed, see above). Finally, participants performed 10 blocks of 64 trials of the task (640 trials in total): the whole session lasted around 80 minutes.

### 2.3. MEG and EEG recording and preprocessing

Simultaneous EEG and MEG data were recorded with a sampling rate of 600Hz during task performance. A 275-channel whole-head axial gradiometer system (CTF-275 by VSM Medtech Inc., Vancouver, Canada) was used to record electromagnetic brain activity (0.016–150Hz filter bandwidth and first-order spatial gradient noise cancellation). Head movements were continuously monitored using 3 coils placed at the nasion and the two preauricular points. EEG was recorded continuously from 7 scalp electrodes placed at frontal (Fz, FC1, FC2), central (Cz), and parietal (Pz) sites, and at the two mastoids (TP9, TP10). The reference electrode was placed on the tip of the nose, the ground electrode on the forehead. One bipolar EOG derivation was recorded from 2 electrodes placed on the supra-orbital ridge of the left eye and infra-orbital ridge of the right eye. For each participant, a 3D MRI was obtained using a 3T Siemens Magnetom whole-body scanner (Erlangen, Germany), locations of the nasion and the two preauricular points were marked using fiducials markers. These images were used for reconstruction of individual head shapes to create forward models for the source reconstruction procedures (see part 2.6).

MEG and EEG data were processed offline using the software package for electrophysiological analysis (ELAN Pack) developed at the Lyon Neuroscience Research Center (Aguera et al., 2011).

MEG data was processed as followed: (1) Raw signals were band-stop-filtered between 47 and 53 Hz, 97 and 103 Hz, and 147 and 153 Hz (zero-phase shift Butterworth filter, order 3) to remove power-line artifacts. (2) An independent component analysis (ICA) was performed on MEG signals filtered between 0.1 and 40 Hz. (3) Component topographies and time courses were visually inspected to determine which components were to be removed (eye-movements and heartbeat artifacts) through an ICA inverse transformation. (4) The ICA inverse transformation was applied to the band-stop filtered MEG signals (resulting from step 1), 2 to 5 components were removed in each participant. (5) Trials contaminated with muscular activity or any other remaining artifacts were excluded automatically using a threshold of 2200 femtoTesla (maximum dynamic range allowed for the duration of a trial).

EEG data was processed as followed: (1) It was band-pass filtered between 0.1 and 40 Hz (zero-phase shift Butterworth filter, order 3). (2) Eye artifacts were removed from the EEG signal by applying a linear regression based on the EOG signal, because of the small number of recorded EEG channels which prevented to use ICA. (3) Trials contaminated with muscular activity or any other remaining artifacts were excluded automatically using a threshold of 150 microvolts (maximum dynamic range allowed for the duration of a trial).

Only trials for which the participant had answered correctly were retained. Trials for which the head position differed of more than 10 mm from the median position during the 10 blocks were also excluded from the analyses. For all participants, more than 80 % of trials remained in the analyses after rejection. Finally, both MEG and EEG data were band-pass filtered between 0.2 and 40 Hz (zero-phase shift Butterworth filter, order 3).

### 2.4. Event-related responses in the sensor space

Event-related fields (ERFs) and potentials (ERPs) were obtained by averaging filtered MEG and EEG data locked to each stimulus event: cue-related responses were locked to cue onset, target-related responses were locked to target onset, and distractor-related responses were locked to distractor onset. A baseline correction was applied based on the mean amplitude of the −100 to 0 ms period before the event. To analyze ERFs/ERPs to distracting sounds, for each distractor onset time-range, surrogate distractor ERFs/ERPs were created in the *NoDIS* trials and subtracted from the actual distractor ERFs/ERPs. The obtained distractor ERFs/ERPs were thus free of cue-related activity. Time-courses and topographies of ERFs/ERPs were plotted using ELAN software. Please note that regarding distractor-related responses, only responses to early distracting sounds (*DIS1*) were considered here in order to analyze late components unaffected by target-related responses.

### 2.5 Source localization of event-related fields

Conventional source reconstruction of MEG data was performed using the Statistical Parametric Mapping (SPM12) toolbox (Wellcome Department of Imaging Neuroscience, http://www.fil.ion.ucl.ac.uk/spm). Previously processed ERF data were converted in a SPM-compatible format. Regarding forward modelling, we considered a three-layer realistic Boundary Element Model (BEM), using canonical meshes provided with SPM12 (scalp, inner skull and cortical sheet) and warped to individual MRI to account for each participant anatomy (Mattout et al., 2007). Forward models were computed with the software OpenMEEG (OpenMEEG Software, https://openmeeg.github.io/, (Gramfort et al., 2010). The estimation of sources was subsequently computed separately for each participant using a LORETA method (Pascual-Marqui et al., 2002), as implemented in SPM12. We performed inversions on the time-windows of interest defined using the time-courses of ERFs for each studied event (concatenation of conditions) (see Figure A.1). Regarding cue-related responses, we reconstructed the contingent magnetic variation (CMV, 650 to 1200 ms post-cue onset). Regarding distractor-related responses, we reconstructed the magnetic N1 (N1m, 80 to 130 ms), the magnetic early-P3 (early-P3m, 200 to 250 ms), the magnetic late-P3 (late-P3m, 290 to 340 ms) and the magnetic reorienting negativity (RONm, 350 to 500 ms). Regarding target-related responses, we reconstructed the magnetic N1 (N1m, 70 to 150 ms) and the magnetic P300 (P3m, 250 to 400 ms).

### 2.6. Statistical analyses

#### 2.6.1. Behavioral data

Trials with response before target (false alarm, FA), trials with incorrect responses and trials with no response after target onset and before the next cue onset (miss) were discarded. Percentages of correct responses and median reaction-times (RTs) in the correct trials were computed for each participant and were submitted to three-way repeated-measures ANOVA (rmANOVAs) with CUE category (2 levels: *uninformative, informative*) and DISTRACTOR condition (3 levels: *NoDIS, DIS1, DIS2*) as within-subject factors and GROUP category (2 levels: *controls*, *migraineurs*) as a between-subject factor. Post-hoc comparisons were conducted using t-tests followed by a Bonferroni correction. For all statistical effects involving more than one degree of freedom in the numerator of the F value, the Greenhouse-Geisser correction was applied to correct for possible violations of the sphericity assumption. We report the uncorrected degree of freedom and the corrected probabilities. Statistical analyses were conducted using the software JASP (version 0.9).

#### 2.6.2. ERP – Sensor-level data

For each ERPs, every sample in each electrode within a time-window of interest (650 to 1200 ms for cue-related ERPs, 0 to 650 ms for distractor-related ERPs, and 0 to 500 ms for target-related ERPs) was submitted to a two-way repeated-measures ANOVA (rmANOVAs) with CUE category (2 levels: *uninformative, informative*) as a within-subject factor and GROUP category (2 levels: *controls*, *migraineurs*) as a between-subject factor. Effects were considered significant if p-values remained lower than 0.05 over a 15 ms interval (corresponding to 9 consecutive samples, see Guthrie and Buchwald, 1991).

In case of a GROUP by CUE interaction, post-hoc unpaired t-tests were performed to assess group difference on the ERP difference *informative* minus *uninformative*, for every sample within the time-windows that had been found significant with the rmANOVA. Again, effects were considered significant if p-values remained lower than 0.05 over a 15 ms interval (corresponding to 9 consecutive samples).

#### 2.6.3. ERF - Source-level data

All statistical analyses regarding the activity of cortical sources were conducted using built-in statistical tools in SPM12. To investigate the GROUP and CUE main effects and the CUE by GROUP interaction, a two-way repeated-measure ANOVA was conducted on the value of source activity for each and every cortical vertex. Significance threshold was 0.05 at the cluster level (p-values corrected for family-wise error, cluster forming threshold=0.05). In order to correct for multiple testing (as several time-windows are inspected, see 2.5 above), a subsequent Bonferroni correction has been applied.

## 3. Results

Demographics and results of the HAD, HIT-6 and MIDAS questionnaires are displayed in Table 1. The control and migraine group did not significantly differ in terms of age, education, musical education, anxiety and depression scores (all p>0.3). The control and migraine group did not significantly differ in terms of the pitch difference between the two target sounds (Control & Migraineurs: 1.4 ± 0.2 tones, Controls: 1.4 ± 0.2 tones).

### 3.1. Behavior

Behavioral data are depicted Figure 2. Participants responded correctly in 95,2% of trials. Remaining trials were either incorrect responses (4,3%), false alarms (0,3%) or misses (0,1%).

**Figure 2:**
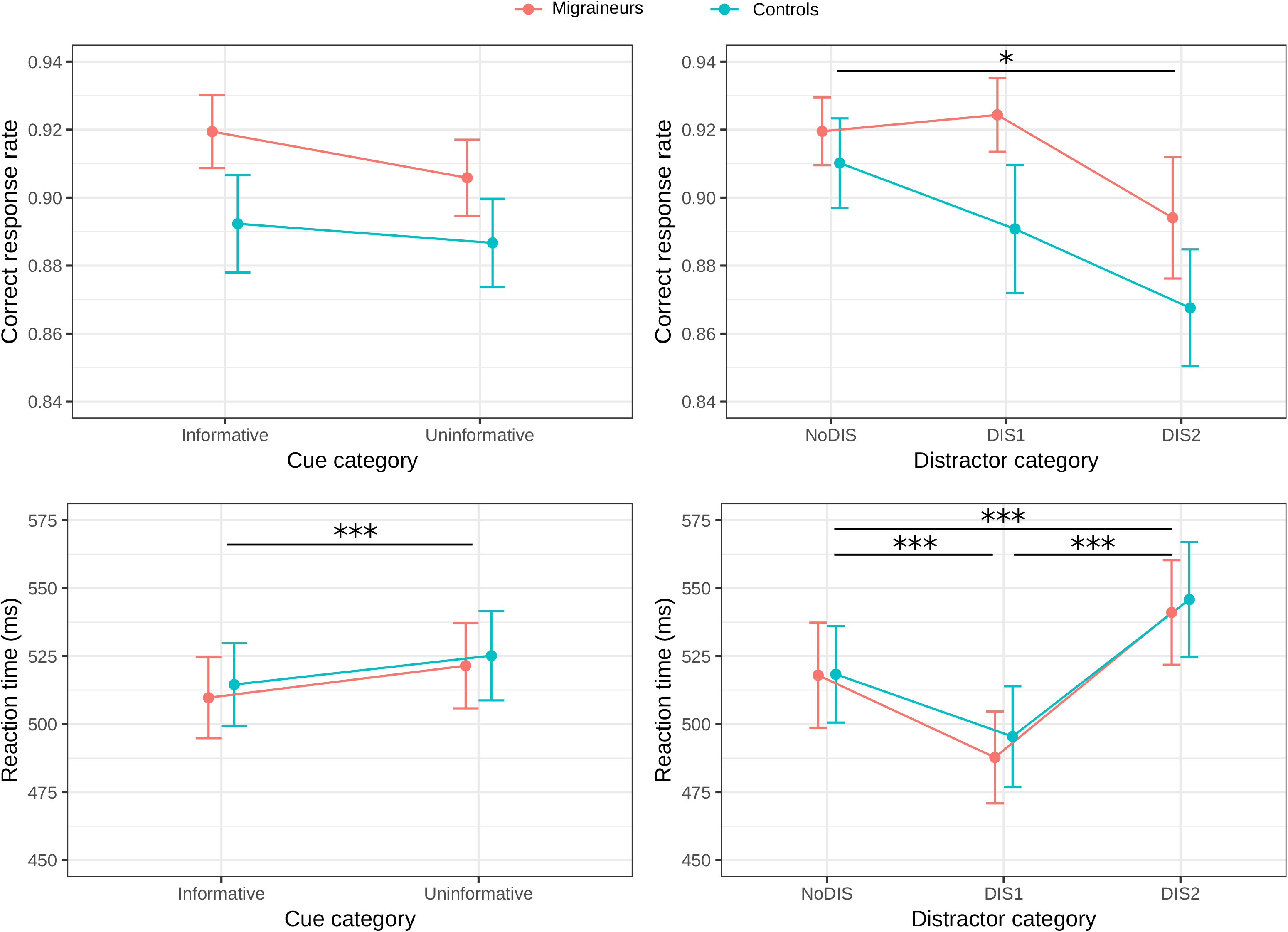
Behavioral results. Mean correct response rate (top) and mean reaction times in milliseconds (bottom) to the target as a function of the GROUP (migraineurs or controls), and as a function of (left) the CUE category (*informative, uninformative*) or (right) the DISTRACTOR category (*NoDIS, DIS1 and DIS2*). ***: p<0.001, *: p<0.05, error bars represent the standard error of the mean.

The two groups did not significantly differ in terms of percentage of correct responses (Migraineurs: 94.2 ± 1,0%, Controls: 95,4 ± 0,7%, F_1,36_=0.92, p=0.34). The percentage of correct responses was not found significantly modulated by the CUE category (F_1,36_=1.8, p=0.18). The DIS category significantly modulated the percentage of correct responses (F_2,72_=4,8, ε=0.99, p=0.011), with a significant decrease in the *DIS2* condition (93,8% ± 0,8%) compared to the *NoDIS* condition (95,5% ± 0.6%, p=0,006), and a marginal decrease compared to the DIS1 condition (95,2% ± 0,7%, p=0,028 – does not resist to Bonferroni correction). No interaction effect was found significant (all p>0.25).

Concerning the median reaction times, both groups did not significantly differ in their performances (Migraineurs: 515 ± 11 ms, Controls: 520 ± 11 ms, F_1,36_=0.013, p=0.91). A significant main effect of CUE (F_1,36_=16.1, p<0.001) was observed with participants responding faster in the *informative* condition than in the *uninformative* condition. A significant main effect of DISTRACTOR (F_2,72_=43.8, ε=0.69, p<0.001) was observed with participants responding faster in trials with an early distracting sound (*DIS1*) (p<0.001) and slower in trials with a late distracting sound (*DIS2*) (p=0.001) compared to trials without distracting sound (*NoDIS*) (for information, DIS1 vs. DIS2, p<0.001). No interaction effect was found significant (all p>0.5).

### 3.2. Event-related responses

#### 3.2.1. Cue-related responses

Regarding source reconstruction, for every time-window of interest, inversions resulted in an explained variance superior to 95% (average across the 38 participants).

In response to visual cues (Figure 3), participants presented occipital ERPs (obligatory visual ERPs) followed by a fronto-central slow negative wave, the contingent negative variation (CNV), which slowly builds up from around 650 ms to 1200 ms post-cue (corresponding to the target onset). The magnetic counterpart of the CNV, the CMV, was visible at the same latencies (Figure A.1). The time-window of interest for subsequent analyses was 650-1200 ms post-cue onset.

**Figure 3:**
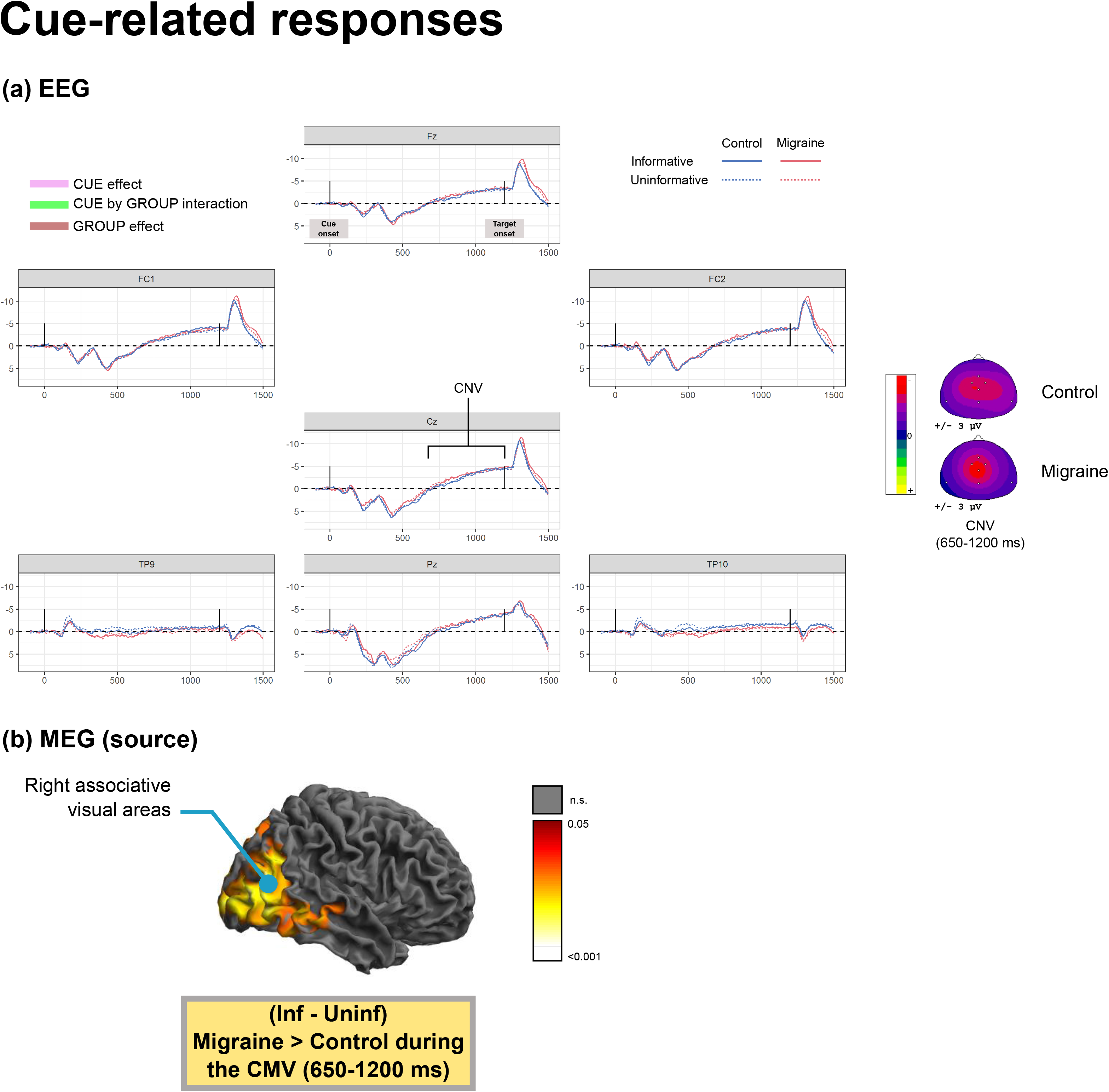
**(a)** Event-related potentials (ERPs) in response to the visual cues as a function of the cue category (informative or uninformative, plain vs. dashed lines) and the group (control or migraine, blue vs. red lines). Time-courses are presented for all EEG sensors. Scalp topographies of the main cue-related responses are presented on the right. The first vertical bar corresponds to the cue onset, the second to the target onset. Statistical analysis of the ERPs during the contingent negative variation (CNV) time-window (650-1200 ms after cue onset) showed no significant effect. **(b)** P-value map (masked for corrected p<0.05, the whiter the more significant) of the pattern of increased cueing effect on brain activation (source-reconstructed MEG data) in the migraine group during the contingent magnetic variation (CMV) time-window (650-1200 ms).

In EEG sensor-level data, neither GROUP nor CUE main effect nor CUE by GROUP interaction were found significant during the time-window of interest.

In MEG source-related data, no GROUP main effect was found significant during the CMV (650-1200 ms). Regarding the CUE main effect, a larger activation of the left occipital, motor and frontal cortices, the bilateral temporo-parietal junctions, and the right parietal and temporal cortices (Brodmann area (BA) 6, 19, 22, 39, 44) was found for *informative* trials compared to *uninformative* trials (Figure A.2). Regarding the GROUP by CUE interaction effect, the effect of the cue information (*informative – uninformative*) was stronger among migraineurs in a cluster including right associative visual areas (BA 7, 19).

#### 3.2.2. Distractor-related responses

In response to distracting sounds (Figure 4), participants presented an expected sequence of ERPs. It includes the fronto-central N1, the fronto-central early-P3 (~270 ms), the fronto-parietal late-P3 (~330 ms) and the frontal reorienting negativity (RON, ~410 ms). The fronto-central N1 comprises two subcomponents: the sensory component of N1 (~95 ms, with polarity inversion at the mastoids) and the orienting component of the N1 (~130 ms, with no polarity inversion at the mastoids). Their magnetic counterparts, respectively labelled in the following as N1m, early-P3m, late-P3m and RONm, were visible at similar latencies (Figure A.1).

**Figure 4:**
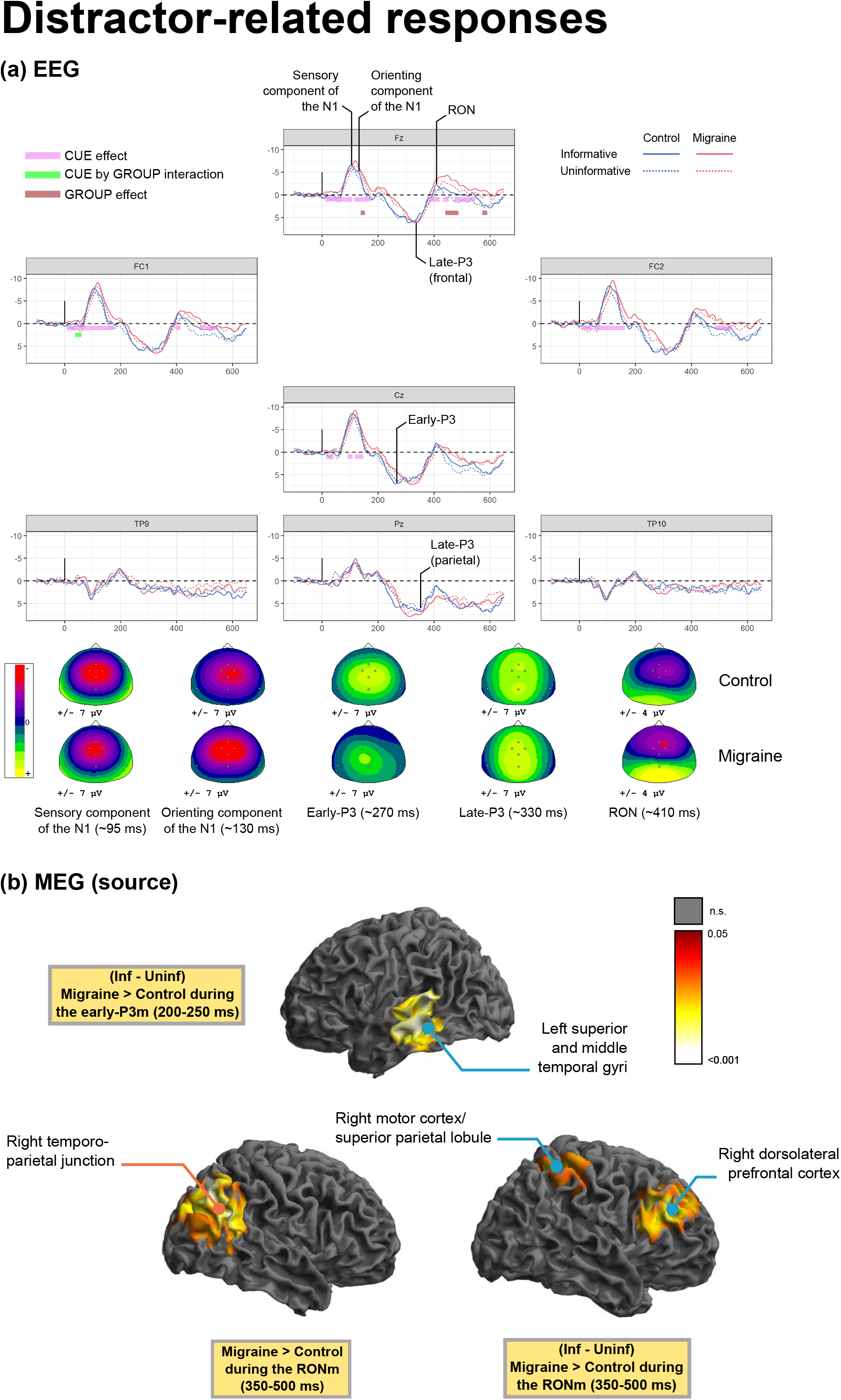
**(a)** Event-related potentials (ERPs) in response to the distracting sounds as a function of the cue category (*informative* or *uninformative*, plain vs. dashed lines) and the group (control or migraine, blue vs. red lines). Time-courses are presented for all EEG sensors. Scalp topographies of the main distractor-related responses are presented below time-courses. GROUP by CUE rmANOVA was applied to ERPs: significant effects (p<0.05 over 15 consecutive ms) correspond to the colored boxes. **(b)** P-value map (masked for corrected p<0.05, the whiter the more significant) of the pattern of increased brain activation in the migraine group during the magnetic reorienting negativity (RONm) time-window (350-500 ms) and the patterns of increased cueing effect on brain activation in the migraine group during the early-P3m (200-250 ms) and the RONm time-windows.

In EEG data, the orienting component of the N1 (138-153 ms) and the RON (440-487 ms then 572-590 ms) were found significantly larger in migraineurs than in controls at Fz. A non-significant trend towards a decreased early-P3 in migraine could be observed. The GROUP by CUE interaction was significant on FC1 in the P50 latency range, prior to the N1 (38-60 ms). Post-hoc analyses confirmed that migraineurs show an increased cueing effect (*informative – uninformative*) during those latencies, with a more positive deflection in *uninformative* trials compared to the control group. Regarding the CUE main effect, during the first 150 ms and during the RON from 380 to 550 ms, responses were found significantly more negative in *informative* trials than in *uninformative* trials at fronto-central electrodes.

In MEG source-related data, at the latencies of the early-P3m (200-250 ms), migraineurs presented an increased cueing effect (*informative – uninformative*) in the left superior and middle temporal gyri (BA 21, 22). At the latencies of the RONm (350-500 ms), migraineurs presented a greater activation of the right angular gyrus (BA 39) which is part of the right temporo-parietal junction (rTPJ), and an increased cueing effect (*informative – uninformative*) in the right dorsolateral prefrontal cortex (BA 9), right frontal eyes fields (BA 8) and right superior parietal lobule and motor cortex (BA 4, 7).

#### 3.2.3. Target-related responses

In response to target sounds (Figures 5, 6), in terms of ERPs, participants presented a fronto-central N1 composed of the sensory component of N1 (~95 ms) and the orienting component of the N1 (~130 ms), followed by a parietal P300 (after 250 ms). Their magnetic counterparts, respectively labelled in the following as N1m and P3m, were observed at similar latencies (Figure A.1).

**Figure 5:**
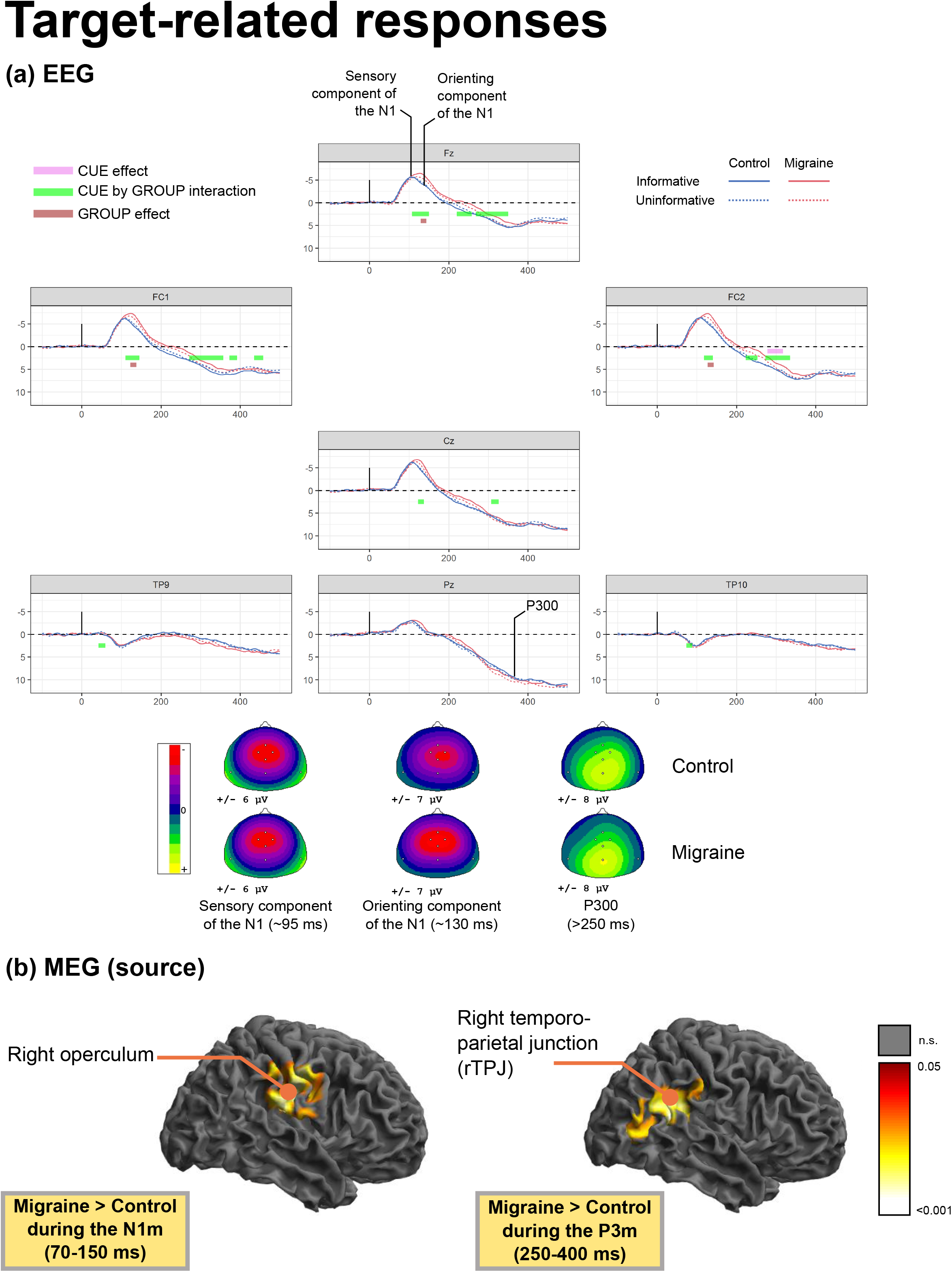
**(a)** Event-related potentials (ERPs) in response to the target sounds as a function of the cue category (informative or uninformative, plain vs. dashed lines) and the group (control or migraine, blue vs. red lines). All EEG sensors are presented. Scalp topographies of the main target-related responses are presented below time-courses. GROUP by CUE rmANOVA was applied to ERPs: significant effects (p<0.05 over 15 consecutive ms) correspond to the colored boxes. **(b)** P-value map (masked for corrected p<0.05, the whiter the more significant) of the pattern of increased brain activation in the migraine group during the N1m and P3m time-window (respectively 70-150 ms and 250-400 ms).

**Figure 6:**
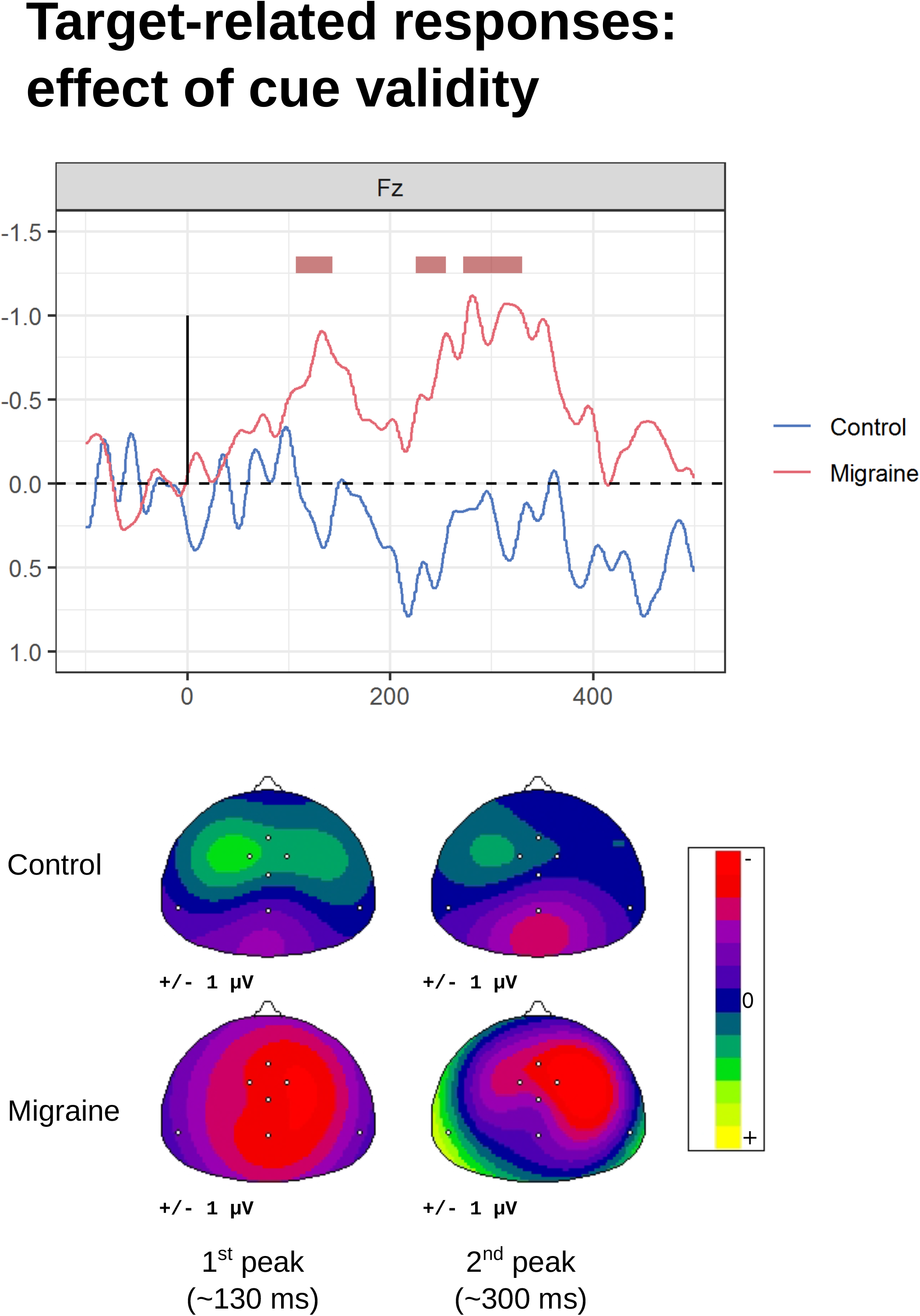
Difference event-related potentials (ERPs) in response to the target (*informative* minus *uninformative* trials), only Fz is presented here. Significant group effects (p<0.05 over 15 consecutive ms) correspond to the brown boxes. Please note the two peaks of the negative difference (Nd) present in the migraine group but absent for the control group.

In EEG data, the orienting component of the N1 on frontal electrodes (Fz, FC1, and FC2) was found larger in migraineurs than in controls (around 130 ms). The GROUP by CUE interaction was significant on fronto-central electrodes around 125 ms and 300 ms (with a significant CUE main effect between 278 and 317 ms at FC2). Difference ERPs (*informative – uninformative*, see Figure 6) showed that contrary to controls participants, migraineurs displayed a frontal negative wave (Negative difference, Nd) comprising two mains peaks (~130 ms and ~300 ms). Post-hoc analyses showed that these two negatives peaks were significantly more negative among migraineurs on frontal electrodes (Fz, FC1, and FC2).

In MEG source-related data, at the latencies of the N1m (70-150 ms), migraineurs presented a larger activation of the right operculum (BA 40). At the latencies of the P3m (250-400 ms), migraineurs presented a larger activation of the right TPJ. Moreover, at the same latencies, a larger activation of the right frontal cortex (BA 9, 47) and of a cluster comprising the right angular gyrus and right occipital gyri (BA 7, 39) was found significant in uninformative trials compared to informative trials (Figure A.2).

## 4. Discussion

Attention in migraine was investigated here using complementary methods. Behavioral data provided us three independent measures of top-down attention, bottom-up attention and phasic arousal in both the patient and control groups. Event-related potentials and fields provided complementary information on brain dynamics as EEG is more sensitive to radial and deep sources while MEG is more sensitive to tangential sources (Ahlfors et al., 2010). Event-related potentials (ERPs) helped to identify precisely which attentional process is potentially dysfunctional in migraine. MEG data, thanks to its superior spatial resolution (Hämäläinen et al., 1993), helped through source reconstruction to pinpoint some of the cortical correlates underlying those alterations. If no behavioral differences were observable between migraineurs and healthy participants, migraine was here associated with elevated responses following distracting sounds (orienting component of the N1 and Re-Orienting Negativity, RON) and following target sounds (orienting component of the N1), conjoined with an increased recruitment of the right temporo-parietal junction. In addition, migraineurs presented an increased effect of the cue informational value on target processing resulting in the elicitation of a negative difference (Nd).

### 4.1. Exacerbated bottom-up attentional effects in migraine

In both participant groups, distracting sounds had opposite behavioral effects depending on the distractor-target interval. Early distracting sounds (*DIS1*) decreased reaction times compared to the condition without distractor (*NoDIS*). This facilitation effect has been previously interpreted as an increase in phasic arousal which improves readiness to respond to any incoming stimulus (Bidet-Caulet et al., 2015; Masson and Bidet-Caulet, 2019). However, late distracting sounds (*DIS2*) resulted in a deterioration of performances (increase of reaction times) compared to early distracting sounds (*DIS1*). This has been previously interpreted as the transient effect of attentional capture by the distracting sound (Bidet-Caulet et al., 2015; Masson and Bidet-Caulet, 2019).

There is no observable evidence that the attentional capture and arousal effects of the distracting sounds were different among migraineurs compared to control participants at the behavioral level. This result is in line with a previous study finding no increased impact over performance of visual distractors during a visual cueing task in migraine (Mickleborough et al., 2016).

However, at the cortical level, migraineurs presented an increased orienting component of the N1 to distracting sounds while the sensory component remained unaltered. The orienting component of the N1 corresponds to the orienting component III described by Näätänen and Picton (Näätänen and Picton, 1987) and is only elicited by infrequent stimuli (Alcaini et al., 1994b). It follows the obligatory sensory component of the N1 and it is considered to be linked to the orienting response to unexpected incoming stimuli (Alcaini et al., 1994b). Increased N1 has been previously reported in migraine interictally (Sable et al., 2017) and also specifically its orienting component (Demarquay et al., 2011; Morlet et al., 2014). These results suggest that the orienting response to distractors is increased in migraine. Unaltered sensory component of the N1 (or earlier responses such as the P50) to the distractor or the target sound argues against an early dysfunctional sensory gating in migraine.

The reorienting negativity (RON) was also increased among migraineurs. The RON is considered to reflect the reorienting of attention towards task-relevant stimuli after distraction (Munka and Berti, 2006; Erich Schröger and Wolff, 1998) but the exact cognitive function of this response is still a matter of debate (Horváth et al., 2008). Source reconstruction of MEG data during the RONm time-window revealed an increased activation of the right temporo-parietal junction (rTPJ) in migraineurs. The rTPJ is part of the ventral attentional network considered to be implicated in stimulus-driven attentional control (for a review, see Corbetta and Shulman, 2002) and is activated by salient unexpected sounds (Salmi et al., 2009). Therefore, enhanced rTPJ activation could reflect an exacerbated bottom-up attentional capture by the distracting sounds in migraine. The rTPJ has also been proposed to play a crucial role in both voluntary and involuntary shifts of attention (Corbetta et al., 2008). In this line, its increased recruitment could also be the necessary consequence of a disproportionate orienting response towards the distracting sound which calls for a more powerful reorientation process towards the task.

Migraineurs also presented an increased orienting component of the N1 to target sounds compared to control participants. Target sounds appear to induce strong orientation responses in migraineurs despite their predictability and low salience. This is consistent with a previous auditory oddball study which reported increased orienting component of the N1 in migraine even for standard sounds (Morlet et al., 2014). Moreover, an increased activation of the rTPJ in migraine could be observed during the P300m time-window, confirming the exacerbation of the orienting response towards target sounds among migraineurs.

These results suggest that migraineurs present an increased orienting response towards both expected relevant and unexpected irrelevant sounds, indicating exacerbated bottom-up attentional processes in migraine. This effect would be mediated, at least in part, by the increased recruitment of the rTPJ, a major node of the ventral attention network (Corbetta and Shulman, 2002). Using fMRI, atypical activation during a visual task (Mickleborough et al., 2016) and functional connectivity profile (Lisicki et al., 2018b, 2018a) of the rTPJ were found in migraine.

### 4.2. Increased top-down attentional effects in migraineurs

Participants responded faster when the visual cue was informative of the auditory target location, in agreement with previous studies using the Competitive Attention Task (Bidet-Caulet et al., 2015; ElShafei et al., 2018a). This effect has been considered to reflect enhanced anticipatory attention.

The effect of the cue informational value on reaction times was not significantly different between the migraine and the control groups, suggesting no difference in top-down attention at the behavioral level in migraine using this paradigm. To our knowledge, three publications have investigated top-down attention in migraine using visual cueing tasks. None of them observed that migraineurs had a greater top-down attentional enhancement in valid cue trials, which is consistent with our results (Mickleborough et al., 2016, 2011a, 2011b). However, at the cortical level, differences in top-down attentional processes were observed between control participants and migraineurs. During target-related responses, the migraineurs presented a frontal slow negative wave in *informative* trials compared to *uninformative* trials, unlike control participants. This resembles the negative difference (Nd), also referred to as the processing negativity (PN). The Nd has been associated with the active selection of relevant information (Alcaini et al., 1994a; Giard et al., 2000; Näätänen, 1982), suggesting enhanced recruitment of voluntary attention in migraineurs.

Moreover, the effect of the cue information was found more pronounced among migraineurs in visual association areas during the CMV preceding targets and in temporal areas during the early-P3m to distracting sounds. Interestingly, a similar effect was found during the RONm to distractors in the dorsolateral prefrontal cortex and the superior parietal lobule, two major nodes of the dorsal attentional network implicated in voluntary top-down attention (Corbetta et al., 2000; Corbetta and Shulman, 2002).

However, no clear evidence of an increased CNV/CMV in migraine could be found using this paradigm. The CNV reflects both attentional anticipation and motor preparation to an imperative stimulus (for a review on the CNV, see Brunia and van Boxtel, 2001), for the CMV, see Elbert et al., 1994; Gómez et al., 2004). These results are inconsistent with previous studies which considered that a wider CNV is a clinical marker of migraine (Kropp et al., 2015; Kropp and Gerber, 1995, 1993; Schoenen and Timsit-Berthier, 1993), which correlates with disease duration (Kropp et al., 2015, 2000) and fails to habituate (Kropp et al., 2015; Siniatchkin et al., 2003). This discrepancy could result from differences in the methods. Previous studies used a simple protocol with a warning signal and an imperative stimulus, separated by a 3-second inter-stimulus interval (while we used here only a one second delay), and the tasks only required motor preparation (while here also attentional processes were at play during the anticipation period).

These results suggest that migraineurs engaged more top-down attentional processes during target processing and anticipation, but also during distractor processing, compared to control participants.

### 4.3. Attention dysfunction in migraine

We hypothesized that migraine is associated with exacerbated bottom-up and/or deficient top-down attention processes, resulting in increased responsiveness to irrelevant information. In consideration of the present data, the reality appears more complex than our hypothesis:

1. Increased brain responses to target and distracting sounds do suggest that the orienting response to attended and unattended sounds is exacerbated in migraine. This is quite consistent with anecdotal reports from migraineurs where they mention being easily distracted by their environment (Sacks, 1992). Migraineurs report higher self-perceived levels of attention difficulty than healthy controls (Carpenet et al., 2019; Lévêque et al., 2020). It is noteworthy that there exists a comorbidity of migraine with attention deficit and hyperactivity disorder (ADHD) (Fasmer et al., 2012; Paolino et al., 2015; Salem et al., 2017).
2. However, at the behavioral level, contrary to our hypothesis, distracting sounds did not have a more pronounced effect on performance in migraine, nor did *informative* cues have a weaker effect in migraineurs. Literature about cognition and attention in migraine is quite contrasted. Neuropsychological evaluations of migraine patients in the literature did not report any major cognitive impairment during the interictal period (Gil-Gouveia et al., 2016; Pearson et al., 2006) but some psychometric tests have linked migraine with diverse minor cognitive alterations (Annovazzi et al., 2004; Calandre et al., 2002; Hooker and Raskin, 1986; Mongini et al., 2005; Zeitlin and Oddy, 1984). Attention in general has been investigated in migraine using specific psychometric tests (for a review, see Vuralli et al., 2018): some studies did not find any interictal attentional alterations in adults (Burker et al., 1989; Conlon and Humphreys, 2001; Koppen et al., 2011); while others have reported moderate impairment of attention during the interictal state (Mulder et al., 1999; Pellegrino Baena et al., 2018; Pira et al., 2000). Conflicting findings in the literature about attention in migraine could be explained by (a) the wide range of psychometrics tests used in the aforementioned studies suggesting that the precise cognitive and attentional processes investigated may vary from study to study, (b) the magnitude of attentional alterations in migraine might be small to moderate.
3. Finally, top-down effects were found increased in migraine as evidenced by event-related potentials and source reconstruction. To our knowledge, increased top-down attentional effects have never been reported in past articles, whether these consisted in behavioral or neuroimaging studies. During attention tasks, migraineurs show either worse or equal performances compared to healthy participants (Vuralli et al., 2018). The present results do not necessarily suggest that migraineurs have superior, more effective top-down attentional mechanisms: they more likely reflect that, in the context of our task, migraineurs have voluntarily engaged more attentional resources in order to be task-efficient.

A good balance between top-down and bottom-up attention is essential to remain task-efficient while still being aware of one’s own environment. The stronger involvement of top-down attentional functions may be seen as a compensatory strategy that migraineurs have developed to cope with heightened bottom-up orienting responses for each and every incoming sound. An increased recruitment of top-down attention would maintain the top-down/bottom-up balance at an operational state, preventing any behavioral impairment. However, it is likely that maintaining such an equilibrium in migraine would be costlier in terms of cognitive resources.

What are the implications of such attentional dysfunctions to the pathophysiology of migraine, and especially to sensory symptoms? The association between attentional difficulties and interictal hypersensitivity in migraine has been validated by a recent questionnaire study from our lab (Lévêque et al., 2020). Several explanations might account for the observed relationship between attention difficulties and sensory hypersensitivity in migraine. Lévêque and colleagues (2020) proposed three hypotheses to explain this relationship. (1) Sensory hypersensitivity would be caused, at least partially, by attentional difficulties linked to migraine: increased bottom-up attention in migraine could lead to sensory overload, as inputs from the environment trigger an orienting response regardless of their actual relevance. (2) Attentional difficulties may be caused by an increased sensitivity to environmental stimuli: sensory amplification associated to migraine would exacerbate attention capture by external stimulation and therefore would produce attention difficulties in the everyday life. (3) Both hypersensitivity and attention alterations would emerge from one’s predisposition to develop migraine: neurochemical imbalances at the core of the migraine pathophysiology might be the source of both dysfunctions. Future studies should aim at exploring the causal links between attention, cognitive load and hypersensitivity in migraine, at cortical and sub-cortical levels. Finally, the knowledge that migraine is associated to disturbed attentional processes may help to shape future recommendation towards migraineurs for the management of their sensory symptoms.

## Acknowledgments

The acquisition of electrophysiological and imaging data was performed at the CERMEP imaging center, we thank Sebastien Daligault and Claude Delpuech from the MEG department for their technical assistance. This work was supported by the French National Research Agency (ANR) Grant ANR-14-CE30-0001-01 (to Aurélie Bidet-Caulet and Anne Caclin). This work was performed within the framework of the LABEX CORTEX (ANR-11-LABX-0042) and the LABEX CeLyA (ANR-10-LABX-0060) of Université de Lyon, within the program “Investissements d’Avenir” (ANR-16-IDEX-0005) operated by the French ANR. The study was approved by the local ethical committee (Comité de Protection des Personnes SUD EST III) and registered under the ID number NCT02791997.

## Supplementary figures

**Figure A.1:**
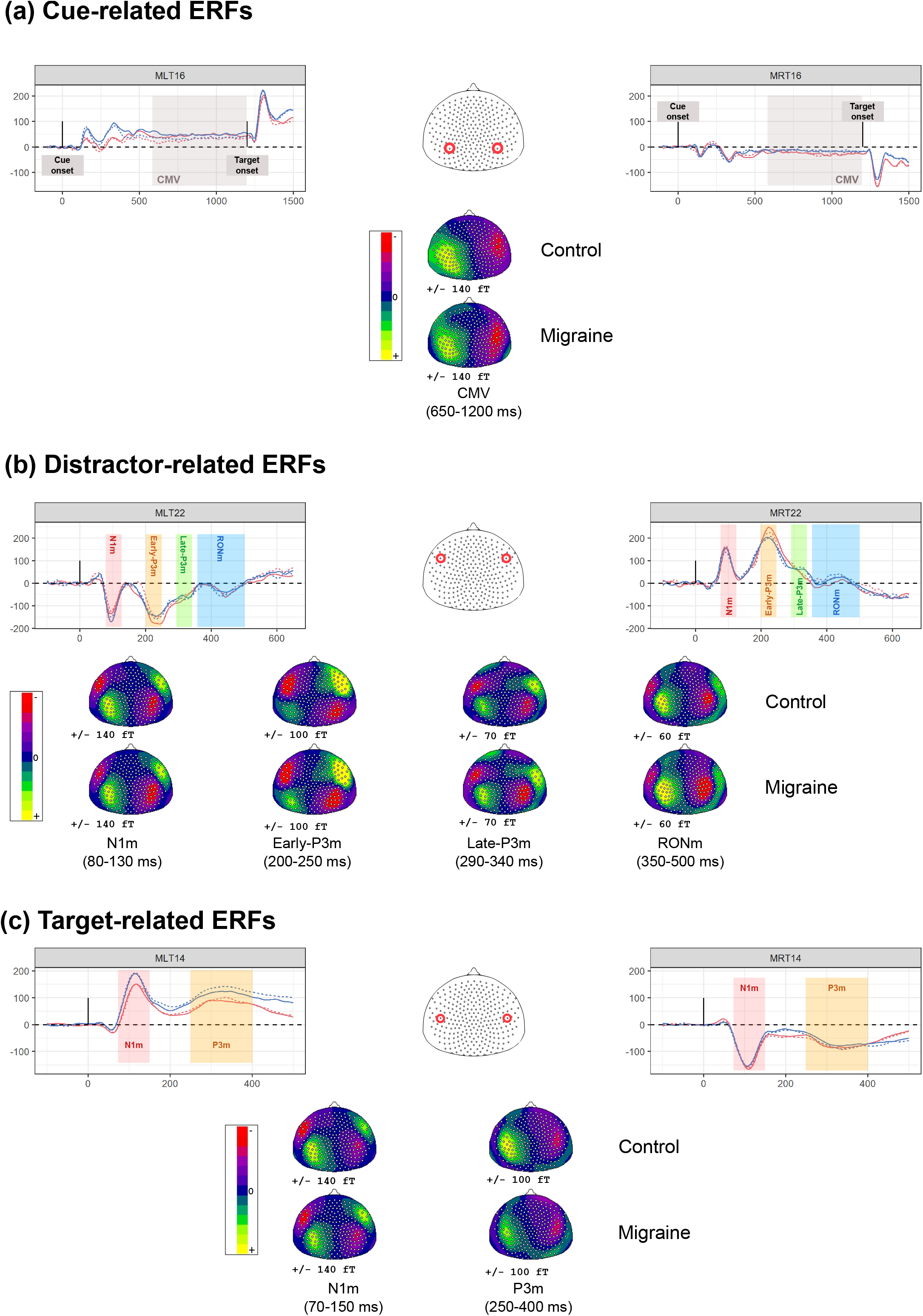
ERFs in response to the cue **(a)**, the distracting sound **(b)** and the target **(c)** as a function of the cue category (informative or uninformative, plain vs. dashed lines) and the group (control or migraine, blue vs. red lines). Time-courses are presented for two selected MEG sensors. Scalp topographies of the main event-related responses are presented under the time-courses: time-windows chosen for scalp topographies matched those used for source reconstruction of MEG data in the article. For all events, the first vertical bar corresponds to the onset of the stimulus; for the cue, the second vertical bar corresponds to the target onset.

**Figure A.2:**
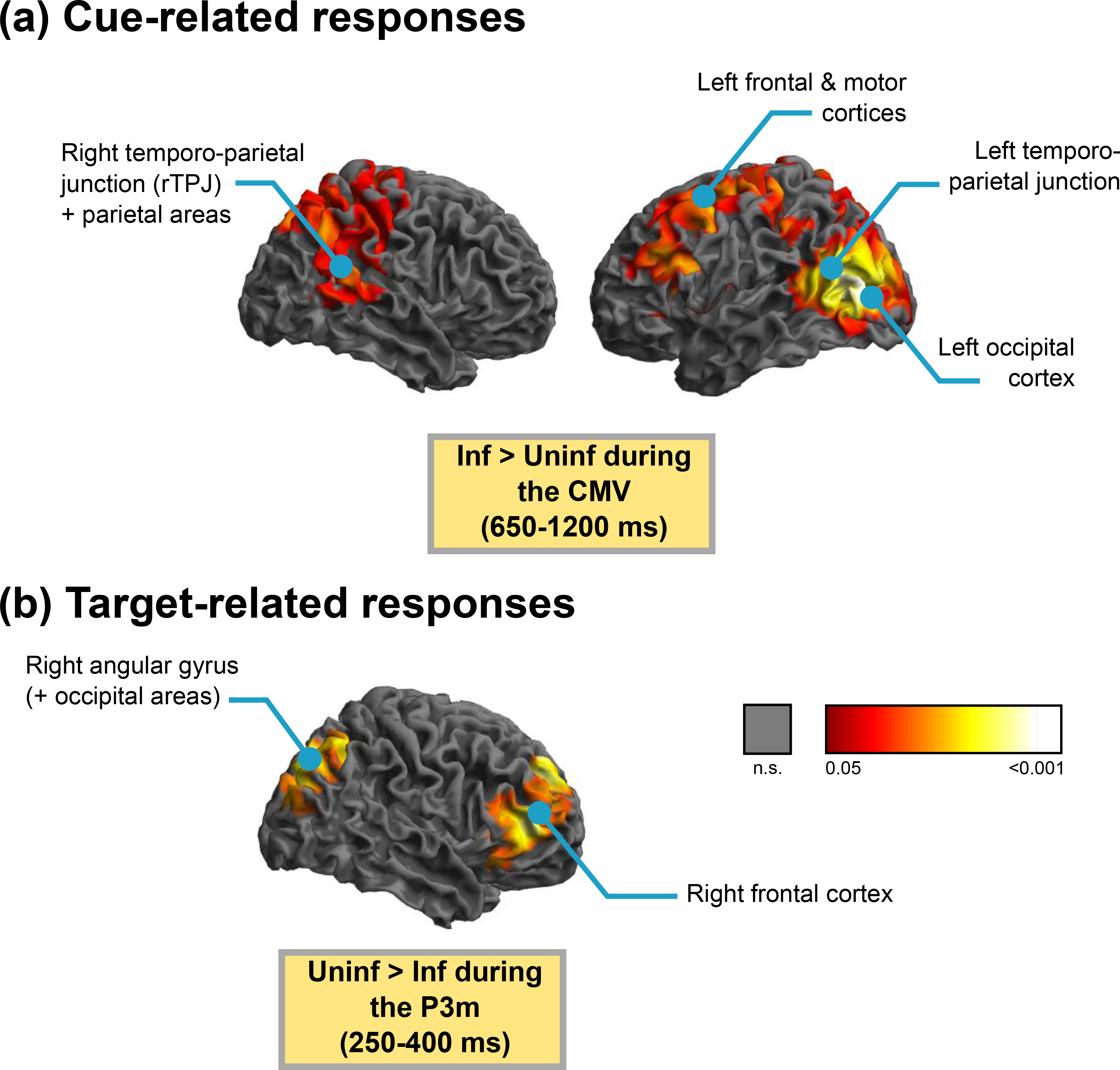
P-value map (masked for corrected p<0.05, the whiter the more significant) of the pattern of increased brain activation **(a)** in informative trials during the CMV in response to the cue (650-1200 ms) and **(b)** in uninformative trials during the P3m in response to the target (250-400 ms).

## Footnotes

1 Pitch discrimination is required in the task described below, and is1 an ability increasing with musical practice.

